# An MHV macrodomain mutant predicted to lack ADP-ribose binding activity is severely attenuated, indicating multiple roles for the macrodomain in coronavirus replication

**DOI:** 10.1101/2021.03.30.437796

**Authors:** Lynden S. Voth, Joseph J. O’Connor, Catherine M. Kerr, Ethan Doerger, Nancy Schwarting, Parker Sperstad, David K. Johnson, Anthony R. Fehr

## Abstract

All coronaviruses (CoVs) contain a macrodomain, also termed Mac1, in non-structural protein 3 (nsp3) which binds and hydrolyzes ADP-ribose covalently attached to proteins. Despite several reports demonstrating that Mac1 is a prominent virulence factor, there is still a limited understanding of its cellular roles during infection. Currently, most of the information regarding the role of CoV Mac1 during infection is based on a single point mutant of a highly conserved asparagine-to-alanine mutation, which is known to largely eliminate Mac1 ADP-ribosylhydrolase activity. To determine if Mac1 ADP-ribose binding separately contributes to CoV replication, we compared the replication of a murine hepatitis virus (MHV) Mac1 mutant predicted to dramatically reduce ADP-ribose binding, D1329A, to the previously mentioned asparagine mutant, N1347A. D1329A and N1347A both replicated poorly in bone-marrow derived macrophages (BMDMs), were inhibited by PARP enzymes, and were highly attenuated *in vivo*. However, D1329A was significantly more attenuated than N1347A in all cell lines tested that were susceptible to MHV infection. In addition, D1329A retained some ability to block IFN-β transcript accumulation compared to N1347A, indicating that these two mutants impacted distinct Mac1 functions. Mac1 mutants predicted to eliminate both binding and hydrolysis activities were unrecoverable, suggesting that the combined activities of Mac1 may be essential for MHV replication. We conclude that Mac1 has multiple roles in promoting the replication of MHV, and that these results provide further evidence that Mac1 could be a prominent target for anti-CoV therapeutics.

**IMPORTANCE:** In the wake of the COVID-19 epidemic, there has been a surge to better understand how CoVs replicate, and to identify potential therapeutic targets that could mitigate disease caused by SARS-CoV-2 and other prominent CoVs. The highly conserved macrodomain, also termed Mac1, is a small domain within non-structural protein 3. It has received significant attention as a potential drug target as previous studies demonstrated that it is essential for CoV pathogenesis in multiple animal models of infection. However, the various roles and functions of Mac1 during infection remain largely unknown. Here, utilizing recombinant Mac1 mutant viruses, we have determined that different biochemical functions of Mac1 have distinct roles in the replication of MHV, a model CoV. These results indicate that Mac1 is more important for CoV replication than previously appreciated, and could help guide the development of inhibitory compounds that target unique regions of this protein domain.

## INTRODUCTION

Coronaviruses (CoVs) are of the family Coronaviridae in the Nidovirales order and are responsible for a variety of diseases of both clinical and veterinary significance. These diseases range from potentially lethal human respiratory diseases such as severe acute respiratory syndrome (SARS)-CoV, SARS-CoV-2, and Middle East respiratory syndrome (MERS)-CoV; to mammalian gastrointestinal diseases such as porcine epidemic diarrhea virus (PEDV) and avian respiratory diseases such as infectious bronchitis virus (IBV) (1). There exists few vaccines or broad-spectrum therapeutics to prevent and treat CoV-induced disease. As SARS-CoV-2 continues to be a significant health threat, and as there will likely be further CoV outbreaks in the future, there is an urgent need for a better understanding of the mechanisms used by CoVs to promote their replication and cause severe disease.

Like other members of the Nidovirales order, CoV genetic information is stored as non-segmented, positive-sense RNA ranging in size from 26-32 kb, making them members of the class of RNA viruses containing the largest genomes. These genomes can be broken down into the region encoding structural and accessory proteins, comprising approximately 10 kb at the 3’ end of the genome, and the region encoding non-structural proteins (nsps), consisting of approximately 20 kb at the 5’ end of the genome. Conserved structural proteins include the spike, envelope, membrane, and nucleocapsid proteins. Accessory proteins have important roles in viral pathogenesis, such as antagonism of the type 1 IFN antiviral response, but are not essential for *in vitro* viral replication. The nsps are translated into a long polyprotein and are responsible for the virus’ genomic and sub-genomic RNA synthesis. The nsps perform a variety of functions, and include the polymerase, helicase, 2 proteases, and many others. However, the function of many nsps are still being fully determined or remain completely unknown. The largest nonstructural protein, nsp3, contains several modular domains including a ubiquitin-like domain, an acidic domain, one or two papain-like protease (PLP) domains, multiple transmembrane domains, and one or more macrodomains (1, 2).

Macrodomains are globular protein domains present in many different life forms, including humans, yeast, bacteria, and several families of positive-sense RNA viruses. They have a highly conserved “sandwich” structure that includes several central β-sheets surrounded by 3 α-helices on each side (3, 4). The primary biochemical functions of macrodomains are to bind and hydrolyze ADP-ribose from proteins. There are additional macrodomains (Mac2/Mac3) in some CoVs, including SARS-CoV-2, that do not bind ADP-ribose and instead bind to nucleic acids or cellular proteins to promote virus replication (5-11).

ADP-ribosylation is the posttranslational covalent addition of ADP-ribose to proteins by ADP-ribosyltransferases that utilize NAD^+^ as a substrate. ADP-ribose can be added to proteins as single subunits (mono-ADP-ribosylation or MAR) or as chains of multiple subunits (poly-ADP-ribosylation or PAR). This process is performed intracellularly by poly-ADP-ribose polymerases (PARPs), also known as diphtheria toxin-like ADP-ribosyltransferases (ARTDs). There are 17 human PARPs with PARP-1 being the most well-studied, as it mediates most of the PARylation that occurs in the cell. Much less is known about the MARylating PARPs, though many of them are interferon stimulated genes (ISGs) and some have demonstrated anti-viral activities (12). For example, PARP12-mediated ADP-ribosylation impedes the replication of Zika virus by ADP-ribosylating NS1 and NS3, leading to their proteasomal degradation (13). PARP activity is countered by several dePARylating or deMARylating enzymes, including PARGs (polyADP-ribose glycohydrolase), ARHs (ADP-ribosylhydrolases), and macrodomains (14).

Recombinant viruses mutated at a highly conserved asparagine residue in the primary CoV macrodomain (herein referred to as Mac1) have been engineered for multiple CoVs to understand the role of Mac1 during infection. This residue was targeted because its mutation to alanine had eliminated Mac1 phosphatase activity, was later shown to eliminate ADP-ribosylhydrolase activity, and is completely conserved amongst all enzymatically active macrodomains (15-18). Structurally, it is positioned to provide critical hydrogen bonds with the terminal ribose, positioning the ADP-ribose for hydrolysis (15, 19, 20). Recombinant viruses with this mutation generally replicate normally in tissue culture cells, but are highly attenuated *in vivo* (16, 21-23). These reports have established Mac1 as a prominent virulence factor and potential therapeutic target (4). For instance, the SARS-CoV Mac1 mutant virus, N1040A, replicates poorly in mice and induces an increased IFN and pro-inflammatory cytokine response in mice and Calu-3 cells, a bronchial epithelial cell line. (16). These results demonstrated that Mac1 is required for the ability of SARS-CoV to fully inhibit IFN and cytokine induction. Similar results were observed following infection of bone-marrow derived macrophages (BMDMs) with the JHM strain of MHV (JHMV) containing the same asparagine-to-alanine mutation (N1347A) in Mac1 (19, 23). We also found that treatment with PARP inhibitors or siRNAs targeting PARP12 or PARP14 enhanced the replication of N1347A. These results demonstrated that the CoV nsp3 Mac1 domain is required to prevent PARP-mediated inhibition of virus replication (23).

While these studies have provided significant insight into the role of the CoV Mac1 domain, they are largely based on a single asparagine-to-alanine mutation. As this mutation may more significantly impact enzymatic activity than ADP-ribose binding, it remains unclear whether the ADP-ribose binding ability of Mac1 may have additional roles in CoV replication. ADP-ribose can be covalently bound to a number of different amino acids, including acidic, basic, serine, and cysteine residues, while macrodomains have only been shown to hydrolyze ADP-ribose attached to acidic residues. Previous studies on the alphavirus macrodomains showed that mutations separating the ADP-ribose binding and hydrolase activities result in distinct phenotypes during virus infection (24, 25). These results indicate that macrodomain binding to ADP-ribosylated proteins with ADP-ribose attached at non-cleavable residues, such as serine, may have functions distinct from its ADP-ribosylhydrolase activity.

Here, we created recombinant JHMV with two distinct mutations that are predicted to impact Mac1 ADP-ribose binding. We found that a D1329A mutant, predicted to abrogate ADP-ribose binding, had more severe replication defects in cell culture than N1347A but mostly retained its ability to block IFN production when compared to N1347A. PARP inhibitors enhanced the replication of D1329A, and NAD enhancing compounds further decreased its replication, indicating that the defects of this virus was due to PARP activity. Finally, we failed to recover a recombinant virus containing both the D1329A and N1437A mutants, or a separate mutant predicted to have diminished ADP-ribose binding and hydrolysis activity, G1439V, suggesting that the combined activities of Mac1 are essential for JHMV replication.

## RESULTS

### Structure of the JHMV Mac1 ADP-ribose binding pocket

As there is no published structure for the MHV Mac1, we used computer modeling to predict its overall structure and the structure of its ADP-ribose binding pocket (Fig. 1A-B). Not surprisingly, the structure of the MHV Mac1 protein is similar to other CoV Mac1 proteins (Fig. 1C-D). One residue predicted to impact ADP-ribose binding is a highly conserved aspartic acid residue located in a loop region between β2 and α1 of MHV Mac1 (D22) (Fig. 1A-B, E-F). This residue is known to either make a critical hydrogen bond with the N6 nitrogen of the adenine ring or mediate water contacts with this molecule (Fig. 1B) (15, 20, 26, 27). Its position in the ADP-ribose binding pocket is well conserved and largely superimposes with that of the SARS-CoV-2 and MERS-CoV Mac1 proteins (Fig. 1E-F). Mutation of this aspartic acid in multiple macrodomains virtually eliminates ADP-ribose binding, but maintains some ADP-ribosylhydrolase activity (16, 28, 29). Therefore, determining the role of this residue in virus replication and pathogenesis may provide unique insight into the functions of Mac1. Another residue of interest is an asparagine located in a loop between β7 and α6 of MHV Mac1 (N156) (Fig. 1A-B, E-F). This residue is mostly conserved amongst β-CoVs, and in our MHV-1 modeled structure it appears to provide hydrogen bonds to the proximal ribose. It has also been proposed to provide similar interactions in the SARS-CoV and MERS-CoV Mac1 proteins (20, 26). This residue is often found as a hydrophobic amino acid in other CoV and viral macrodomains, including phenylalanine in SARS-CoV-2 Mac1 (30). These side chains appear to be in close contact with the adenine base and may stack against the adenine ring to create water-mediated hydrogen bonds with the proximal ribose (31). Based on these observations, we hypothesized that alanine mutations at these residues would provide further insight into the role of Mac1 ADP-ribose binding in CoV replication and pathogenesis.

**Figure 1.**
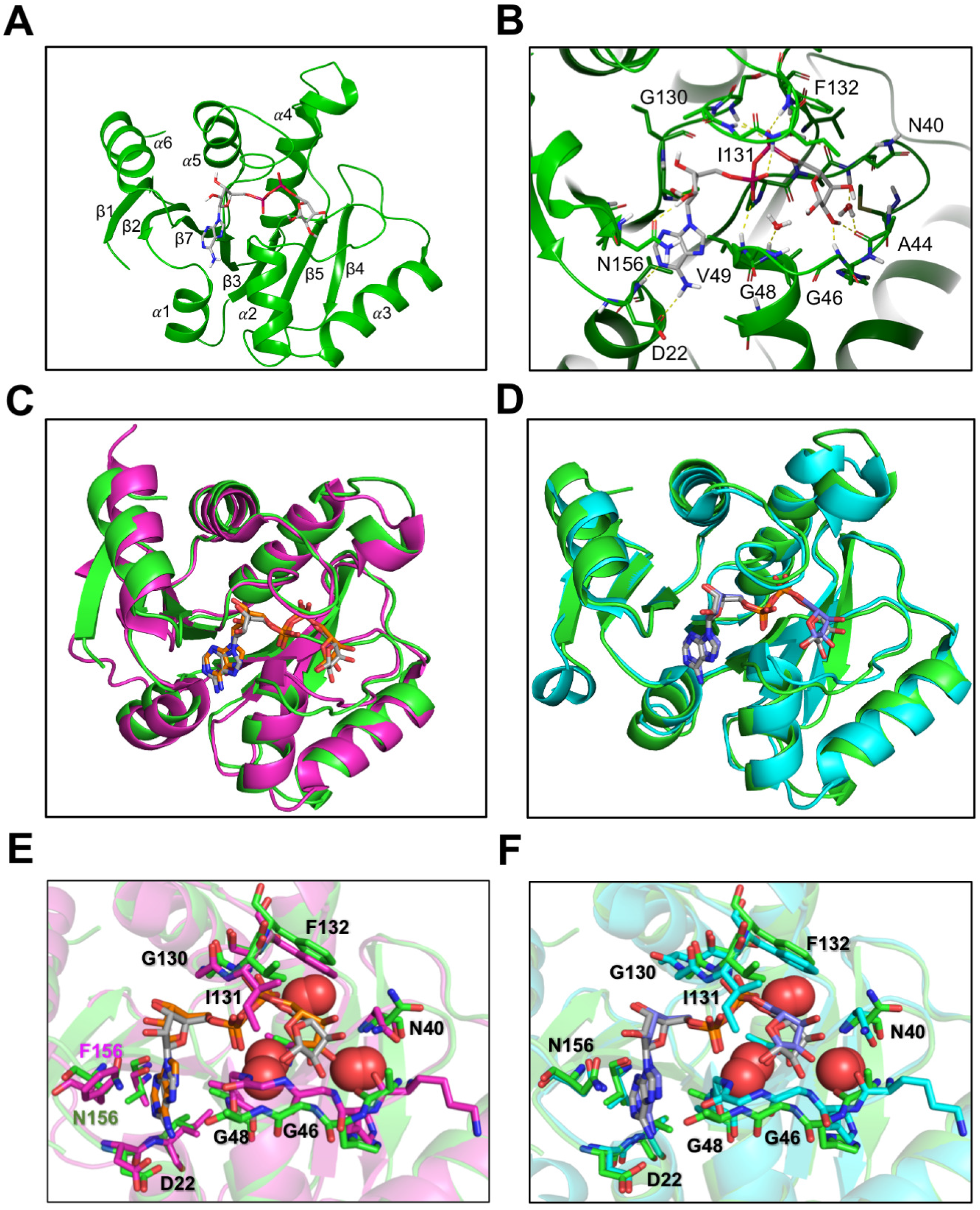
Rosetta-derived structure of the MHV Mac1 protein. **A)** Cartoon representation of MHV Mac-1 with ADP-ribose as determined by Rosetta. **B)** Hydrogen bond interactions (dashed lines) between ADP-ribose and amino acids with modeled water molecules. **C-D)** Superposition of MHV Mac1 (green) (6WOJ) with other CoV Mac1 structures. **C)** SARS-CoV-2 Mac1 with ADP-ribose (magenta) (6WOJ) and **D)** MERS-CoV Mac1 with ADP-ribose (cyan) (5DUS). **E-F)** Superposition of MHV Mac1 (green) with other CoV Mac1 structures highlighting the ADP-ribose binding site. **E)** SARS-CoV-2 (magenta), **F)** MERS-CoV (cyan). The ADP-ribose molecules are colored gray for MHV **A-F)** and are rendered as orange cylinders for SARS-CoV-2 Mac1 **C,E)** and blue cylinders for MERS-CoV Mac1 **D,F)**. Conserved waters are shown as red spheres.

### The MHV Mac1 D1329A virus is highly attenuated in all cell types tested, while N1465A replicates like WT virus

To test the role of the D22 (D1329 in pp1a) and N156 (N1465 in pp1a) residues in the context of MHV replication, we first engineered recombinant JHMV BACs containing the D1329A and N1465A mutations using a two-step Red recombination with the endonuclease I-SceI as previously described (22, 32). These recombinant viruses, termed D1329A and N1465A, were easily recovered and replicated in cell culture. Following reconstitution of virus, we sequenced the Mac1 region and confirmed that these mutations had been retained following passaging.

We first compared the replication of D1329A and N1465A with that of WT and N1347A viruses in BMDMs. We previously showed that N1347A had significantly decreased replication in BMDMs, and initially hypothesized that D1329A and N1465A would be similarly affected. While WT and N1465A viruses were able to replicate to similar levels at all time points, N1347A and D1329A were highly attenuated, showing a greater than 1-log replication defect throughout the infection (Fig. 2A). We next tested the replication of D1329A and N1465A on 17Cl-1 fibroblasts, a common cell line that is highly permissive for MHV replication. As expected, N1347A replicated like WT virus in these cells, as did N1465A, but surprisingly D1329A replicated poorly in these cells with a replication defect of ∼1-log (Fig. 2B-C) and produced reduced levels of viral nucleocapsid (N) and spike (S) protein (Fig. 2D). To confirm that this defect was not due to a second-site mutation in the BAC, we repaired this mutation to create the BAC clone, *rep*D1329. The virus recovered from this BAC clone replicated well and produced viral proteins at WT virus levels in 17Cl-1 cells, confirming that the replication defect was due to mutation of D1329 (Fig. 2C-D).

**Figure 2.**
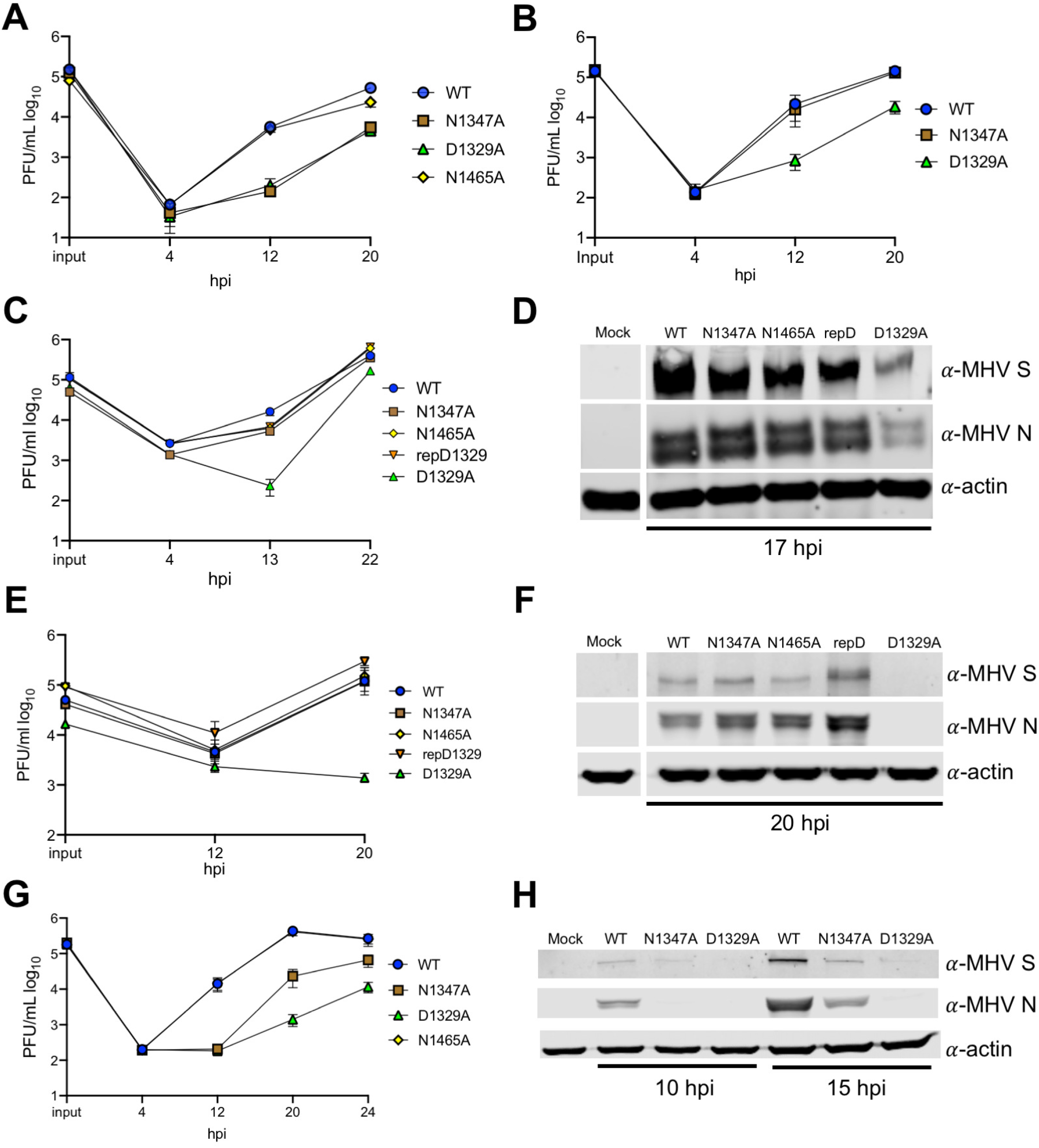
D1329A, but not N1465A, is highly attenuated in both primary cells and in cell lines. BMDM (A), 17Cl-1 (B-D), L929 (E-F), and DBT (G-H) cells were infected with WT, N1347A, D1329A, N1465A, and *rep*D1329 viruses as described in Materials & Methods. Progeny virus was collected at indicated time points and virus titers were determined by plaque assay. In addition, cell lysates were collected and viral protein levels were determined by immunoblotting. The data show one experiment representative of at least two independent experiments (A-C,E,G) with n=4 (A) or n=3 (B,C,E,G) in each experiment. The data in (D,F,H) show one experiment representative of two independent experiments.

Next, we tested whether D1329A was defective in additional MHV susceptible cells. In both L929 fibroblasts and DBTs (delayed brain tumor), an astrocytoma cell line, D1329A was highly attenuated with replication defects of approximately 2-logs, and very little N and S protein accumulated in both cell types compared to WT virus (Fig. 2E-H). Interestingly, we found that N1347A also replicated poorly and produced reduced N and S protein in DBT cells, though it replicated better than D1329A (Fig. 2G-H).

### The MHV Mac1 D1329A virus mostly retains the ability to repress IFN production in BMDMs

The dramatic reduction in replication of D1329A could indicate that it had lost all ADP-ribose binding and hydrolase activity, while N1347A maintains at least partial activity for one or both of these functions. Alternatively, D1329A may primarily impact ADP-ribose binding while N1347A primarily affects Mac1 enzymatic activity. Current biochemical data based on analyses of similar macrodomains favor the latter hypothesis. Previous reports have found that the aspartic acid mutation leads to complete or nearly complete loss of binding but these mutants retained some hydrolase activity. In contrast, the asparagine mutation leads to the near complete loss of Mac1 hydrolysis activity but had only a 2 to 3-fold reduction in ADP-ribose binding (16, 25, 28, 29, 33).

To further test these opposing hypotheses, we measured the level of IFN-β mRNA at 6 and 12 hpi in BMDMs infected with WT, N1347A, D1329A, and N1465A. We previously found that N1347A infection led to a >1-log increase in IFN-β transcript levels in BMDMs (23). We predicted that if D1329A had less overall activity compared to N1347A, its infection would lead to an equivalent or greater increase in IFN-β transcript levels than N1347A. However, we might expect the D1329A mutant infected cells to have reduced IFN-β transcript levels compared to N1347A infected cells if hydrolysis was primarily responsible for blocking IFN production. Here, infection with N1347A resulted in a >19-fold increase in IFN-β mRNA levels at 12 hpi and a smaller difference at 6 hpi (Fig. 3A), consistent with our previous report. N1465A-infected cells had similar IFN-β transcript levels to those infected with WT virus, further indicating that this mutation does not affect Mac1 function. However, instead of observing further increases of IFN-β mRNA following D1329A infection, we found that IFN-β mRNA levels in BMDMs were reduced 3.5-fold when compared to N1347A, despite having roughly the same level of viral genomic RNA (gRNA) in the cells in this specific experiment (Fig. 3A-B). These results indicate that D1329A retains some ability to block IFN-β transcription, and thus likely has more, and not less, ADP-ribosylhydrolase activity than N1347A. These results, in combination with results in Fig. 2, suggests that Mac1 utilizes multiple mechanisms to promote virus replication and block innate immune responses in cell culture.

**Figure 3.**
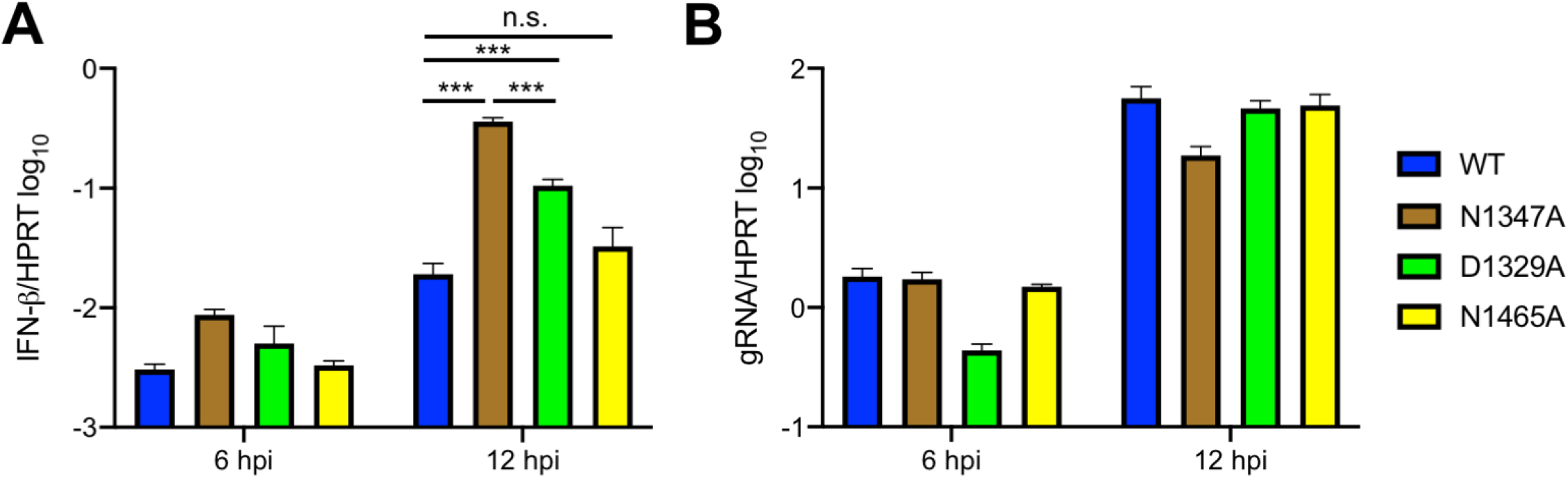
D1329A infection results in lower IFN-β mRNA levels than N1347A infection in BMDMs. (A-B) BMDMs were infected with WT, N1347A, D1329A, or N1465A recombinant virus. Cells were collected at the indicated times post infection and RNA was purified. Grna (A) and IFN-β (B) mRNA levels were determined by RT-qPCR using primers listed in Table 2 and normalized to HPRT mRNA levels. The data in (A and B) show one experiment representative of four independent experiments with n=4 for each experiment.

### PARP inhibitors significantly enhance the replication of MHV D1329A, while boosting the PARP substrate NAD^+^ with nicotinamide riboside further inhibits D1329A replication

The D1329 residue is known to be critical for ADP-ribose binding from multiple studies (28, 29), however it is also conceivable that this mutation could affect other functions of the CoV macrodomain, such as Papain-Like Protease (PLPro) binding (34). To address this possibility, we treated cells with inhibitors that target host PARP enzymes. If the ADP-ribosyltransferase activity of PARP enzymes are restricting the replication of D1329A, we would expect to see at least partial restoration of virus replication in the presence of PARP inhibitors, as we previously showed with N1347A (23). First, we performed MTT assays to confirm that the PARP inhibitors XAV-939 and Olaparib (2281) did not affect the metabolic capacity of BMDMs or 17Cl-1 cells. BMDM and 17Cl-1 cells were treated with XAV-939 and 2281 for 24 hrs and MTT levels were measured. Neither compound resulted in notable metabolic changes at the working concentration of 10 μM (Fig. 4A-B). In virus replication experiments, both compounds significantly enhanced the replication of D1329A in BMDMs and in 17Cl-1 cells, by approximately 5 and 3-fold, respectively, but had no effect on WT virus (Fig. 4C-D). In addition, the level of enhanced replication in BMDMs for D1329A was similar to that of N1347A. These results indicate that PARP-mediated ADP-ribosylation inhibits the replication of D1329A.

**Figure 4.**
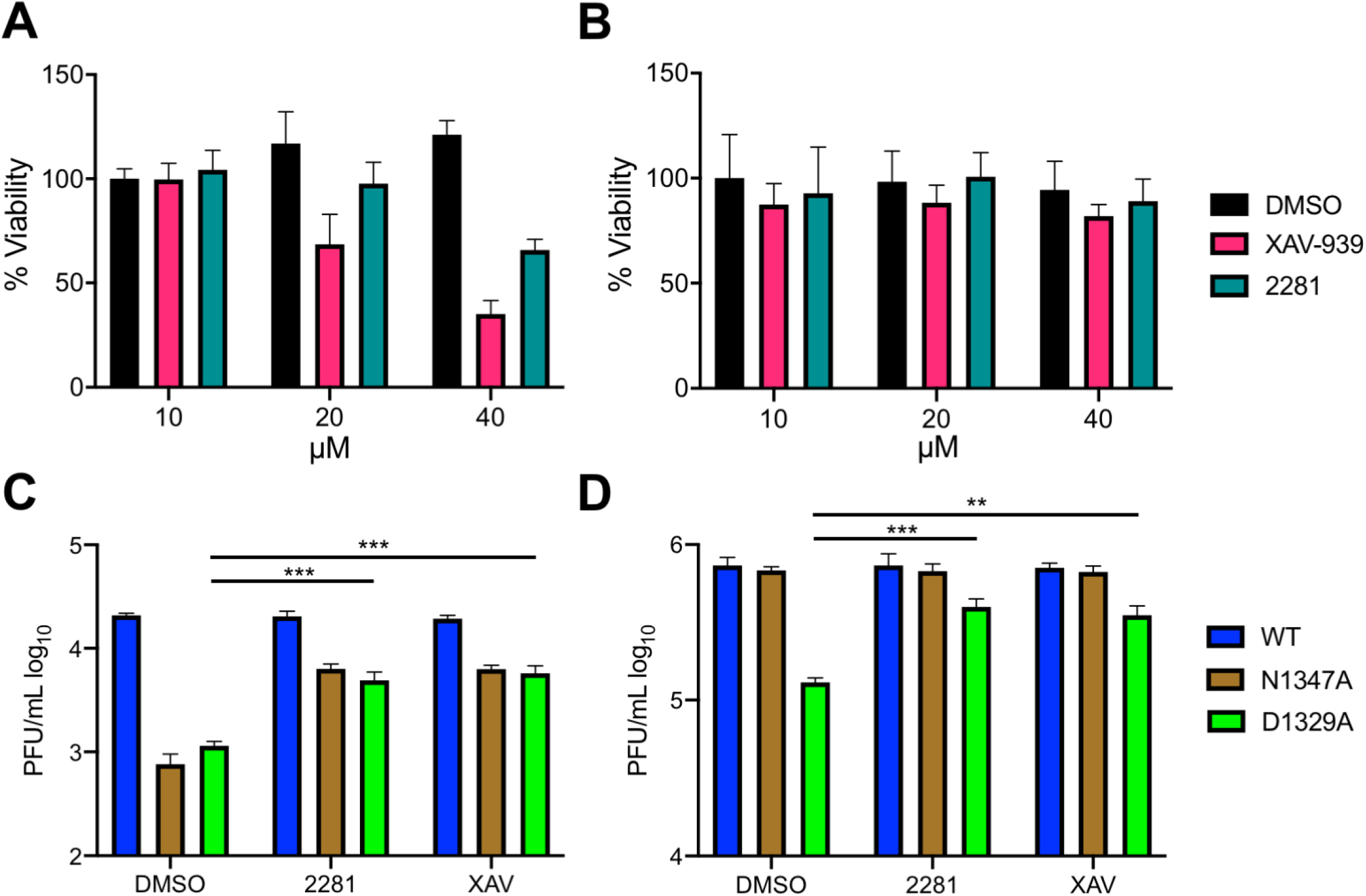
D1329A replication is significantly increased by the addition of PARP inhibitors. (A-B) BMDMs (A) and 17Cl-1 cells (B) were treated with indicated compounds, and at 24 hours, cell viability was measured using an MTT assay as described in Materials & Methods. (C-D) WT BMDMs (C) and 17Cl-1 cells (D) were infected with WT, N1347A, or D1329A, and then treated 0.25% DMSO, 10 μM Olaparib (2281), or 10 μM XAV-939 as described in Materials & Methods. Progeny virus was collected at indicated time points and virus titers were determined by plaque assay. The data in (A-D) show one experiment representative of two independent experiments with n=4 (A,B) or n=3 (C,D) for each experiment.

To provide further evidence for PARP-mediated inhibition of D1329A, we hypothesized that increasing PARP activity using nicotinamide riboside chloride (NR) would lead to further reduction in D1329A replication. NR enhances PARP activity by increasing the intracellular levels of the PARP substrate, NAD^+^ (35, 36). We recently showed that NAD levels are depleted following MHV infection, and that restoring NAD levels with NR and other NAD boosting compounds both increased PARP activity and decreased the replication of N1347A (37). We first confirmed that NR did not significantly decrease the metabolic activity of BMDM or 17Cl-1 cells at and above the working concentration of 100 μM (Fig. 5A-B). Rather than decreasing metabolic activity, 17Cl-1 cells treated with NR seemed to have slightly increased metabolic activity, though this was not statistically significant (Fig. 5B). For the infection, we pretreated these cells with NR for 4 hrs, then infected them with WT and D1329A, added fresh NR after the infection and then collected both cell-free and cell-associated virus at 20 hpi. Similar to our results with N1347A, we found that NR significantly reduced the replication of D1329A virus in both cell types by >5-fold, but had no impact on WT virus (Fig. 5C-D). These results indicate that NR enhanced PARP activity that further reduced D1329A virus replication, but was countered by Mac1 in WT virus infected cells.

**Figure 5.**
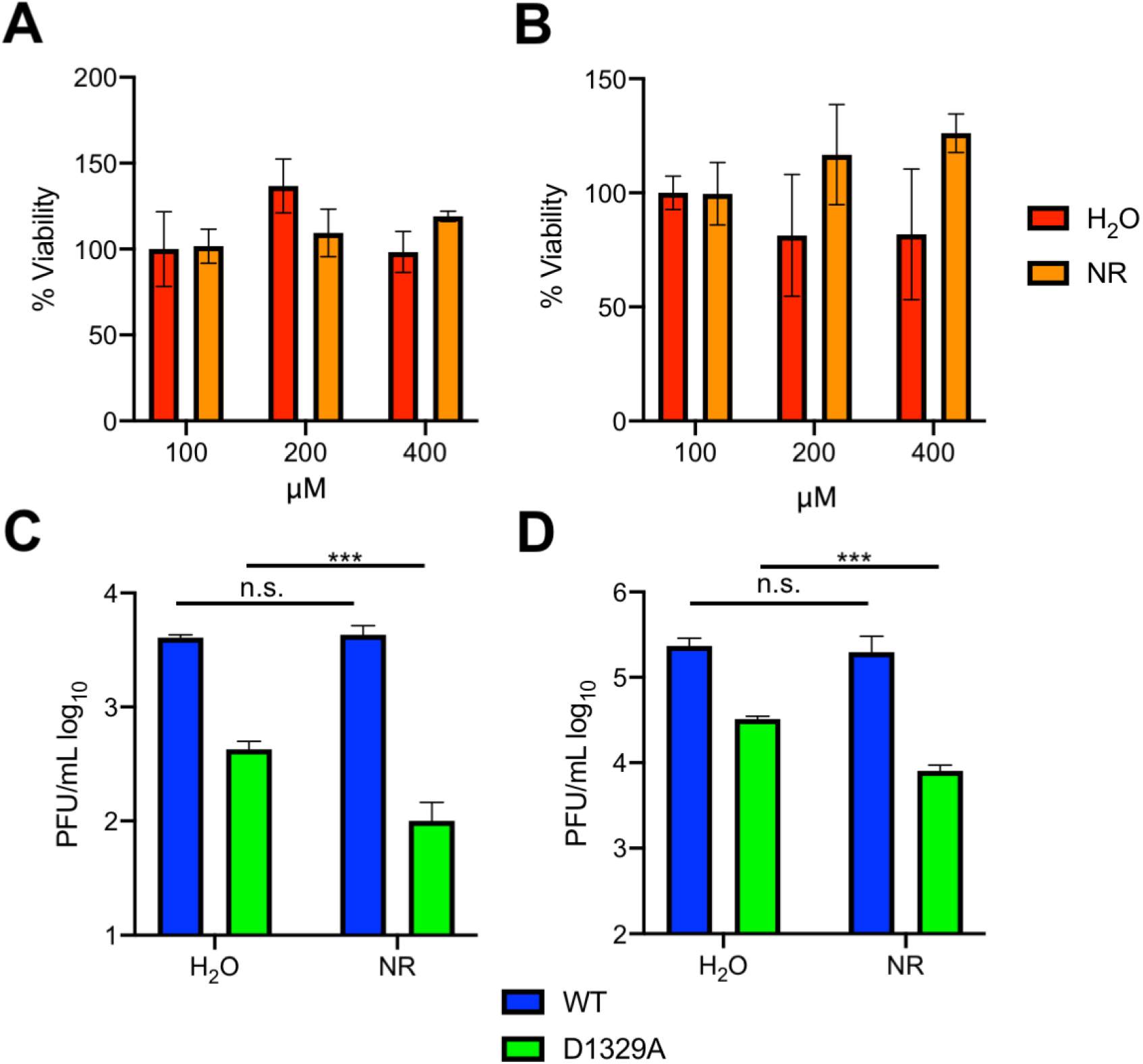
The addition of NR, a precursor of the PARP substrate NAD^+^, further decreases the replication of D1329A. (A-B) BMDMs (A) and 17Cl-1 cells (B) were treated with NR as described above, and at 24 hours cell viability was measured using an MTT assay as described in Methods. (C-D) WT BMDMs (C) and 17Cl-1 cells (D) were either mock treated (H_2_O) or treated with NR as described in Materials & Methods. Progeny virus was collected at indicated time points and virus titers were determined by plaque assay. The data in (A-D) show one experiment representative of two independent experiments with n=4 (A,B) or n=3 (C,D) for each experiment.

### Infection of mice with Mac1 ADP-ribose binding mutants

As the results in cell culture may not mimic what happens *in vivo*, we tested the effect of these mutations in mice, where WT JHMV causes a lethal encephalitis (22, 38). We infected C57BL/6 mice with recombinant WT, N1347A, and N1465A JHMV at 3×10^4^ PFU and monitored weight loss as a clinical sign of disease progression (Fig. 6A-B). N1347A was included as an attenuated control virus as it was previously shown to cause minimal disease in B6 mice (22). We hypothesized that N1465A would cause disease similar to WT virus, as all cell culture experiments indicated that this mutation had not affected Mac1 function. Indeed, N1465A infected mice lost weight and were euthanized due to severe disease at nearly the same rate as WT infected mice, while N1347A did not cause any weight loss and all mice survived its infection (Fig. 6A-B). These results provide further evidence that mutation of N1465 does not significantly affect the function of Mac1.

**Figure 6.**
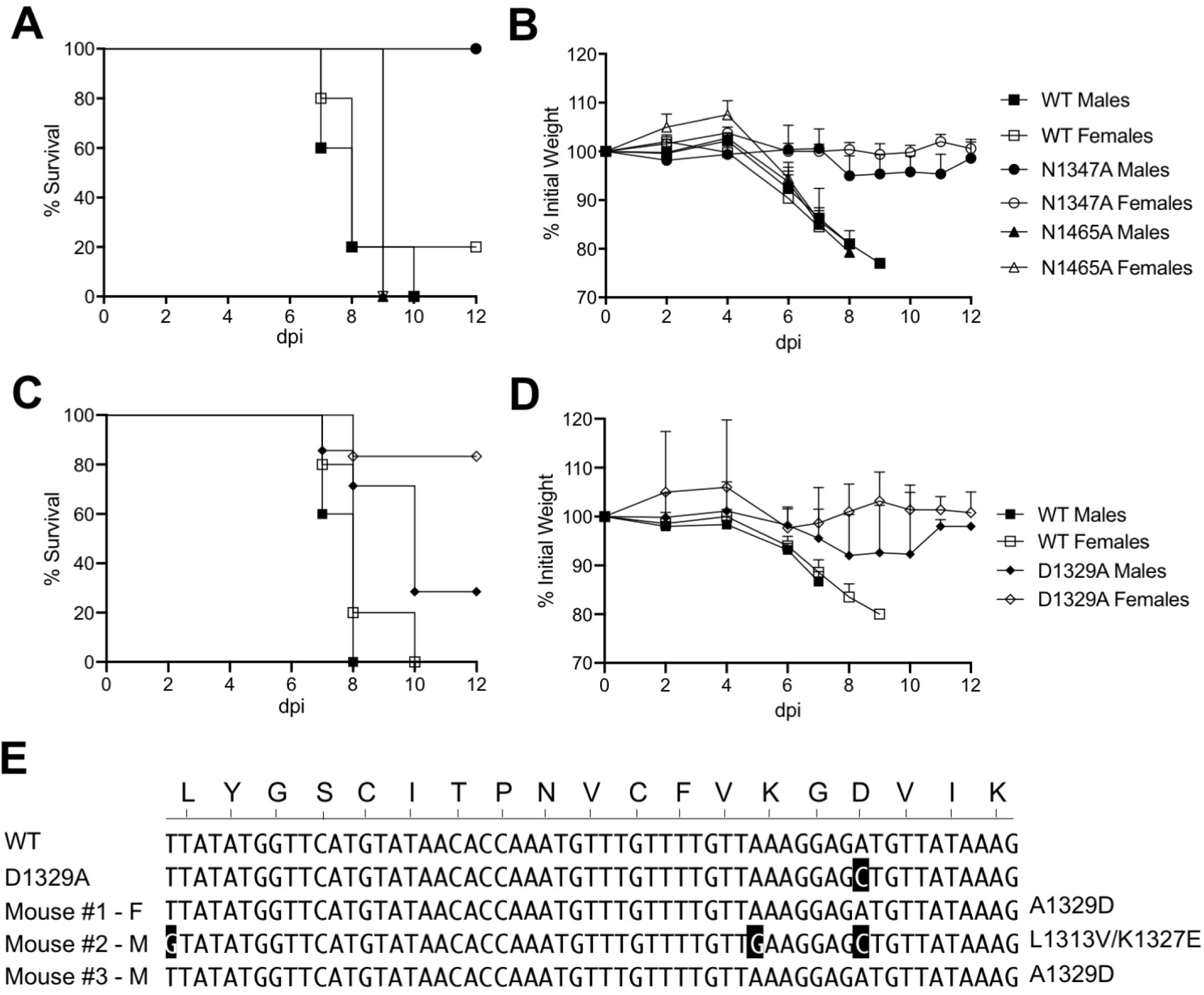
D1329A, but not N1465A, is highly attenuated *in vivo*. (A-B) WT male and female B6 mice were infected with 3×10^4^ PFU of WT, N1347A, and N1465A intranasally and monitored for survival and weight loss daily for 12 days. The data show the combined results of two independent experiments. WT, n=5 for male and female mice; N1347A, n=5 for male and female mice; N1465A, n=4 for male and female mice. (C-D) WT male and female B6 mice were infected with 3×10^3^ PFU of WT and D1329A as described above. The data show the combined results of two independent experiments. WT, n=5 for male and female mice; D1329A, n=7 for male and n=6 for female mice. (E) D1329A virus readily reverts *in vivo*. The brains of 3 mice (2 male (M); 1 female (F)) which succumbed to infection with D1329A were harvested and their viral RNA was amplified by RT-PCR with Mac1 specific primers. The PCR product was sequenced by Sanger sequencing and analyzed using DNA Star software. Mouse #1 and #3 reverted to wild-type virus sequence, while mouse #2 evolved 2 new mutations, L1313V & K1327E.

Next, we infected mice with 3×10^3^ PFU of WT and D1329A virus. This low dose was used because titers of the D1329A virus stocks were significantly lower than other viruses. Regardless, WT virus still caused significant weight loss and the infected mice succumbed to this infection at roughly the same rate as did those receiving the higher dose of virus. Interestingly, the D1329A virus did lead to weight loss and lethality in 5 out of 7 male mice and 1 out of 6 female mice (Fig. 6C-D). Since this result was unexpected, we collected the brains of 1 female and 2 male mice that succumbed to infection with D1329A and sequenced the Mac1 region of the MHV genome. We found that in 2 of the 3 mice, the alanine reverted back to aspartic acid, while in the third mouse there were 2 second-site mutations in the N-terminus of the macrodomain, L1313V and K1327E (Fig. 6E). As all mice sequenced had reverted virus, it appears that D1329A replicates poorly *in vivo* and is especially prone to reversion. We conclude that D1329A is highly attenuated *in vivo*.

### Recombinant JHMV with mutations predicted to impact both ADP-ribose binding and hydrolysis are not recoverable

To further test the hypothesis that Mac1 has multiple roles in virus replication, we created a D1329A/N1347A double mutant BAC clone. We hypothesized that if the D1329 residue confers a unique role in virus replication when compared to N1347, then a double-mutant virus may be even more attenuated than either single mutant. Consistent with this hypothesis, we were unable to recover the D1329A/N1347A virus following 12 transfection attempts from 2 separate BAC clones. Each of these experiments included successful transfections of other BACs as positive controls (Table 1).

**Table 1.** MHV BAC Recovery Rates in BHK-MVR Cells.

| **BAC** | **Virus Recovered/Attempts** | **% Recovery** |
| --- | --- | --- |
| N1347A | 9/9 | 100% |
| D1329A | 9/9 | 100% |
| N1347A/D1329A | 0/12 | 0% |
| G1439V | 4/11 | 36.37% |
| A1438T/G1439V | 6/6 | 100% |

Introducing multiple mutations into the macrodomain could disrupt its structure and thus the inability to recover this virus could be due to impairment of additional nsp3 functions outside of Mac1. To provide further evidence that the loss of Mac1 function is lethal for JHMV, we engineered an additional mutant, G1439V into our JHMV BAC clone. This mutation has previously been introduced into SARS-CoV and SARS-CoV-2 recombinant Mac1 proteins, and both proteins had minimal, if any, ADP-ribosylhydrolase activity (16, 39). Furthermore, computational modeling of an ADP-ribose bound structure of the SARS-CoV and MERS-CoV Mac1 proteins found that ADP-ribose binding was highly unfavorable when introducing the G-V mutation, with a ddG of binding of ∼9 and 10 rosetta energy units (REUs), respectively. After transfection, we only recovered this virus 4/11 times for a 37% recovery rate (Table 1). Upon sequencing the recovered viruses, we found that most had reverted to WT virus. However, one clone instead had evolved a second site mutation of alanine to threonine in the residue immediately upstream of V1439. To determine if this mutation was responsible for the ability of this virus to replicate, we created a A1438T/G1439V mutant BAC clone. We were able to easily recover this virus, indicating that this mutation was the reverting mutation that allowed the outgrowth of one of the G1439V viruses (Table 1). A1438T/G1439V replicated better than D1329A on L929 cells, but had a 3-fold replication defect and produced less viral protein on L929 cells compared to WT virus (Fig. 7A-B). It also did not cause severe disease in mice, indicating this mutation only partially recovered Mac1 functions (Fig. 7C-D). In total, we have created two separate Mac1 recombinant JHMV clones (D1329A/N1347V and G1439V) that were not recoverable without second site or reverting mutations. These results suggest that the combined ADP-ribose binding and hydrolysis activities of Mac1 are essential for JHMV replication.

**Figure 7.**
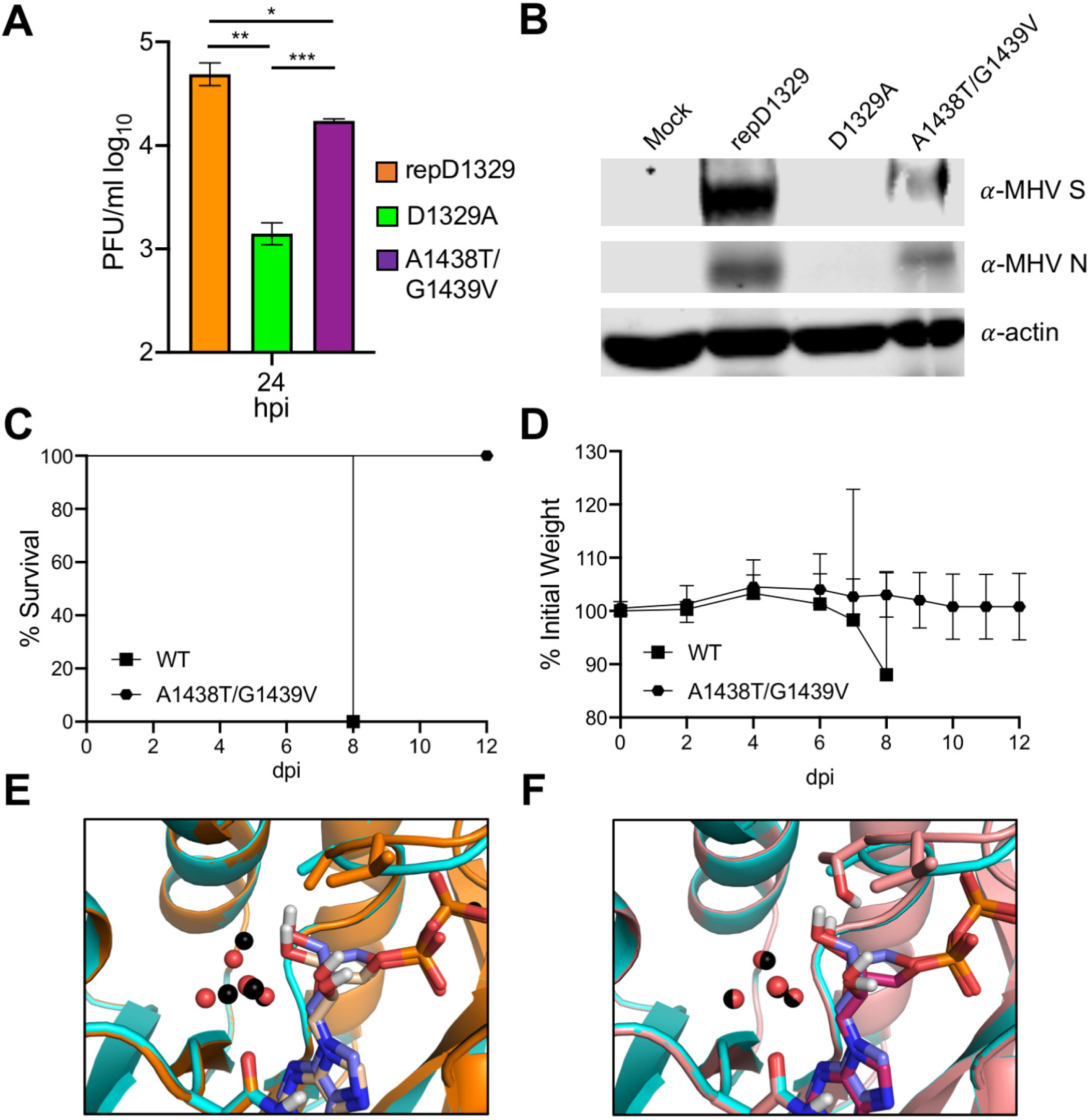
The A1438T/G1439V virus, a revertant of G1439V, is recoverable but replicates at slightly reduced levels compared to WT virus and is attenuated *in vivo*. (A-B) L929 cells were infected with WT, D1329A, and AG1438/1439TV viruses as described in Materials & Methods. Progeny virus was collected at 24 hpi and virus titers were determined by plaque assay. In addition, cell lysates were collected at 24 hpi and viral protein levels were determined by immunoblotting. The data in (A) shows one experiment representative of two independent experiments with n=3 for each experiment. The data in (B) shows one experiment representative of two independent experiments. (C-D) WT male B6 mice were infected with 3×10^3^ PFU of WT and A1438T/G1439V intranasally and monitored for survival (C) and weight loss (D) daily for 12 days. WT, n=3; A1438T/G1439V, n=6. (D-E) Rosetta predicted structures of WT MERS-CoV Mac1 around the distal ribose compared to G128V (E) and A127T/G128V (F) proteins. G128V (orange cylinders) is predicted to cause a disruption of water molecules (E) (WT – red spheres; G128V or A127T/G128V – black spheres). The A127T mutation (salmon cylinders) is predicted to restore this water network back to their original location (F), likely restoring critical hydrogen bonds.

## DISCUSSION

Here we show that the JHMV Mac1 protein domain promotes virus replication in multiple ways, likely by utilizing both its ADP-ribose binding and ADP-ribosylhyrolase activities. Based on prior reports demonstrating the importance of Mac1 for CoV pathogenesis, there has been particular interest in developing therapeutic strategies targeting the SARS-CoV-2 Mac1 domain. (16, 31, 39-44). These results further demonstrate the importance of Mac1 for CoV replication and could have implications in the design of compounds targeting the ADP-ribose binding domain of Mac1.

Previous studies of the alphavirus macrodomain has provided unique insight into the separate binding and enzymatic activities of these protein domains (24, 25, 29, 45). Chikungunya (CHIKV) and Sindbis virus mutations that impact these activities are highly attenuated in cell culture and *in vivo*. In fact, CHIKV macrodomain mutations that reduced hydrolysis activity by as little as 25% are attenuated, and any mutation that eliminated binding activity was unrecoverable (29). In addition, two mutants were further analyzed for their impact on the virus replication cycle, G32S (reduced hydrolysis and binding) and Y114A (decreased hydrolysis but increased binding). While both mutants had reduced virus replication as measured by plaque assay, the G32S mutant had greatly reduced levels of viral RNA and protein production early in the infection, while Y114A had normal levels of viral RNA and protein until later stages of infection. It was concluded that a CHIKV with ADP-ribose binding defects is unable to properly initiate replication, while a hydrolase deficiency negatively affects the later stages of virus replication.

To determine if there may be additional phenotypes associated with the ADP-ribose binding ability of the CoV Mac1, we chose to mutate residues in the adenine binding pocket. First, we targeted N1465, located in a loop between β7 and α6, which appears to provide a hydrogen bond with the proximal ribose (Fig. 1B). Despite its apparent ability to provide interactions with the proximal ribose (20, 26), we found that mutation of this residue to alanine had no impact on JHMV replication in any cell type we tested (Fig. 2), and also did not affect the ability of JHMV to cause disease in mice (Fig. 6A-B). From these results we conclude that N1465 does not contribute significantly to the ability of the JHMV Mac1 to promote replication or cause disease.

Next, we targeted a conserved aspartic acid that makes a critical hydrogen bond with the N7 nitrogen of the adenine base, D1329. While D1329 behaved similarly to N1347A in BMDMs, it was substantially attenuated in multiple cell lines, including 17Cl-1, L929, and DBT cells. Importantly N1347A replicates normally, or with only modest defects in these cells, suggesting that D1329 contributes to virus replication in a unique manner compared to N1347. This is consistent with biochemical data from other macrodomains that demonstrates that the D-A mutation leads to dramatic loss of ADP-ribose binding, while the asparagine mutation only results in a 2-3-fold loss of ADP-ribose binding (25, 28, 29). In mice we found that D1329A often reverted. While in some cases it had reverted to WT, in one case two novel mutations near D1329, L1313V and K1327E appeared. Both residues are found outside of the position of the aspartic acid though not in near contact with the substrate (data not shown), making it unclear if and how either of these amino acid changes helped enhance the replication of D1329A *in vivo*. However, it is possible that the D1329A mutation could have significantly affected the structure in this region of Mac1, causing a rearrangement of these residues. It will be of interest to determine if one or both of these mutations can help restore ADP-ribose binding activity and virus replication of this mutant.

In addition, we have shown that the defect of D1329 is due to PARP-dependent ADP-ribosylation and not an additional Mac1 function. PARP inhibitors increased D1329A replication in 17Cl-1 and BMDMs (Fig. 4), and NR, which increases levels of the PARP substrate NAD^+^, further decreased its replication in these cell types (Fig. 5) (35-37). From these results it is likely that ADP-ribose binding is a critical component of MHV replication. In contrast, D1329A did not induce IFNβ transcript levels to the same degree as N1347A, which indicates that this virus retains some function that is lost with N1347A. A logical explanation is that ADP-ribosylhydrolase activity is responsible for Mac1 inhibition of IFNβ transcript levels, and that N1347A has lost most, if not all of this enzyme activity while D1329A retains at least some hydrolase activity. This is consistent with biochemical data of other CoV Mac1 mutant proteins showing that the N-A mutation largely ablates hydrolase activity while the D-A mutant retains some hydrolase activity (15, 16, 18, 25, 29).

Interestingly, we were unable to recover a double mutant virus (D1329A/N1347A) or a separate mutant designed to negatively impact both binding and hydrolase activities (G1439V). These results indicate that the combined activities of Mac1 may be essential for JHMV replication. Intriguingly, we identified a second site mutation for G1439V in an immediately adjacent residue, A1438T, that allowed it to replicate. Of note, all alphaviruses have a threonine in this position. Based on computer modeling, we predict that this mutation may affect the network of water molecules in the vicinity of these residues and could potentially improve either the ADP-ribose binding or hydrolysis activity of Mac1 (Fig. 7C-D). After recreating an isogenic BAC clone with these two mutations, A1438T/G1439V, we found that this clone was easily recoverable and replicated in cell culture, though it had a mild replication defect compared to WT virus in L929 cells and was attenuated *in vivo*. Regardless, these results confirmed that this mutation allowed us to recover the G1439V mutant virus and provided even more evidence that Mac1 is a potential therapeutic target for CoVs. Mac1 could be targeted by: i) using inhibitors specific to the macrodomain; ii) enhancing PARP activity by increasing NAD^+^ levels by consumption of NR; or iii) a combination of both.

The primary challenge remaining is to identify targets of Mac1 that contribute to the inhibition of Mac1 mutant viruses. Macrodomain proteins have only been demonstrated to remove ADP-ribose from acidic residues (4), and several PARPs that are induced by virus infection ADP-ribosylate acidic residues (29, 46). But recent proteomic analyses have shown that in certain conditions, ADP-ribosylation of acidic residues is rare or even absent (47). Thus, it’s likely that viral macrodomains would also target proteins modified at non-acidic residues such as serine, cysteine, or lysine, even if they cannot hydrolyze these ADP-ribose modifications. Mac1 binding to this modification may dramatically impact the biochemical activities of its target proteins, for instance, by altering protein-protein interactions. While Mac1 is likely to target both cellular and viral proteins during infection, the severe and universal defect of D1329A indicatesthat its binding activity may target an ADP-ribosylated CoV protein. We recently found that the CoV nucleocapsid (N) protein is ADP-ribosylated. Interestingly, this modification was unchanged following infection with N1347A, indicating it is not cleaved by the macrodomain but may be a target for macrodomain binding (48). Regardless, further investigation into how PARP-mediated ADP-ribosylation impacts the function of viral and cellular proteins is likely to uncover unique insights into the replication of CoVs.

## MATERIALS AND METHODS

### Molecular modeling of the MHV Mac1 protein

A model of MHV Mac1 protein bound to ADP-ribose was generated using RosettaCM (49), using the ADP-ribose bound MERS-CoV Mac1 structure, 5DUS, as a template (20). Following the threading step, the pose of ADP-ribose bound to the MERS-CoV macrodomain was included in the refinement process to create a model of the MHV Mac1 protein bound to ADP-ribose. The top scoring model was aligned to the MERS Mac1 protein, and three structural waters that participate in hydrogen bond networks bridging the protein to ADP-ribose (water residues 312, 365, and 384) were added to the MHV model, followed by hydrogen bond optimization and minimization within Maestro by Schrodinger.

### Rosetta Cartesian ddG calculations

The structures of the MERS-CoV and SARS-CoV Mac1 proteins bound to ADP-ribose, 5DUS (20) and 2FAV (15) respectively, were stripped of all waters but the three structural waters bridging the macrodomain to ADP-ribose (20). These structures, as well as the model of the MHV bound to ADP-ribose and the three waters, were prepared for ddG prediction by relaxing into cartesian space using coordinate restraints via Rosetta (50). The lowest energy structure was selected for ddG prediction of the G1439V mutant using Rosetta’s cartesian ddG protocol (51). By default, cartesian ddG protocol runs for three iterations for the wild type and G1439V proteins, and the predicted ddG was taken as the difference between the wildtype and the mutant for the average of the three simulations. Commands are available upon request.

### Cell culture

17Cl-1, Delayed brain tumor (DBT), 17Cl-1, L929, HeLa cells expressing the MHV receptor carcinoembryonic antigen-related cell adhesion molecule 1 (CEACAM1) (HeLa-MHVR), and baby hamster kidney cells expressing CEACAM1 (BHK-MVR) (all cell lines gifts provided by Stanley Perlman, University of Iowa) were grown in Dulbecco’s Modified Eagle Medium (DMEM) supplemented with 10% fetal bovine serum (FBS), 100 U/ml penicillin and 100 mg/ml streptomycin, HEPES, sodium pyruvate, non-essential amino acids, and L-glutamine. Bone marrow-derived macrophages (BMDMs) sourced from WT mice were differentiated by incubating cells in Roswell Park Memorial Institute (RPMI) media supplemented with 10% L929 cell supernatants, 10% FBS, sodium pyruvate, 100 U/ml penicillin and 100 mg/ml streptomycin, and L-glutamine in for seven days. Cells were washed and replaced with fresh media every day after the 4^th^ day.

### Cell viability assay

BMDMs and 17Cl-1 cells were treated with the indicated compounds for 24 hours. Cellular metabolic activity was assessed using a Vybrant MTT Cell Proliferation Assay (Thermo Fisher Scientific) following manufacturer’s instructions.

### Mice

Pathogen-free C57BL/6 (B6) mice were originally purchased from Jackson Laboratories. Mice were bred and maintained in the animal care facility at the University of Kansas as approved by the University of Kansas Institutional Animal Care and Use Committee (IACUC) following guidelines set forth in the Guide for the Care and Use of Laboratory Animals.

### Generation of recombinant pBAC-JHMV constructs

All recombinant pBAC-JHMV constructs were created using Red recombination (primers listed in Table 2). Recombinant WT (rJ^IA^-GFP*rev*N1347) and N1347A (rJ^IA^-GFP-N1347A) MHV were previously described. Recombinant BACs with the D1329A, G1439V, A1438T, and N1465A point mutations in the nsp3 macrodomain were engineered using the Kan^r^-I-SceI marker cassette for dual positive and negative selection as previously described, using the primers listed in Table 2 (32). BAC DNA from Cml^r^ Kan^s^ colonies was analyzed by restriction enzyme digest, PCR, and direct sequencing for isolation of correct clones. PCR and sequencing were done using the following primers located just outside of the Mac1 gene sequence: F 5’-ggctgttgtggatggcaagca-3’ and R 5’-gctttggtaccagcaacggag-3’. Wild-type repaired BACs were engineered by reintroducing the wild-type sequence into the BAC clones containing the D1329A mutation using the same procedure as described above. The resulting BAC clones were termed pBAC-JHMV^IA^-GFP-D1329A, pBAC-JHMV^IA^-GFP-N1465A, pBAC-JHMV^IA^-GFP-D1329A/N1347A, pBAC-JHMV^IA^-GFP-G1439V, pBAC-JHMV^IA^-GFP-G1439V/A1438T, and pBAC-JHMV^IA^-GFP-*rep*D1329.

**Table 2.**
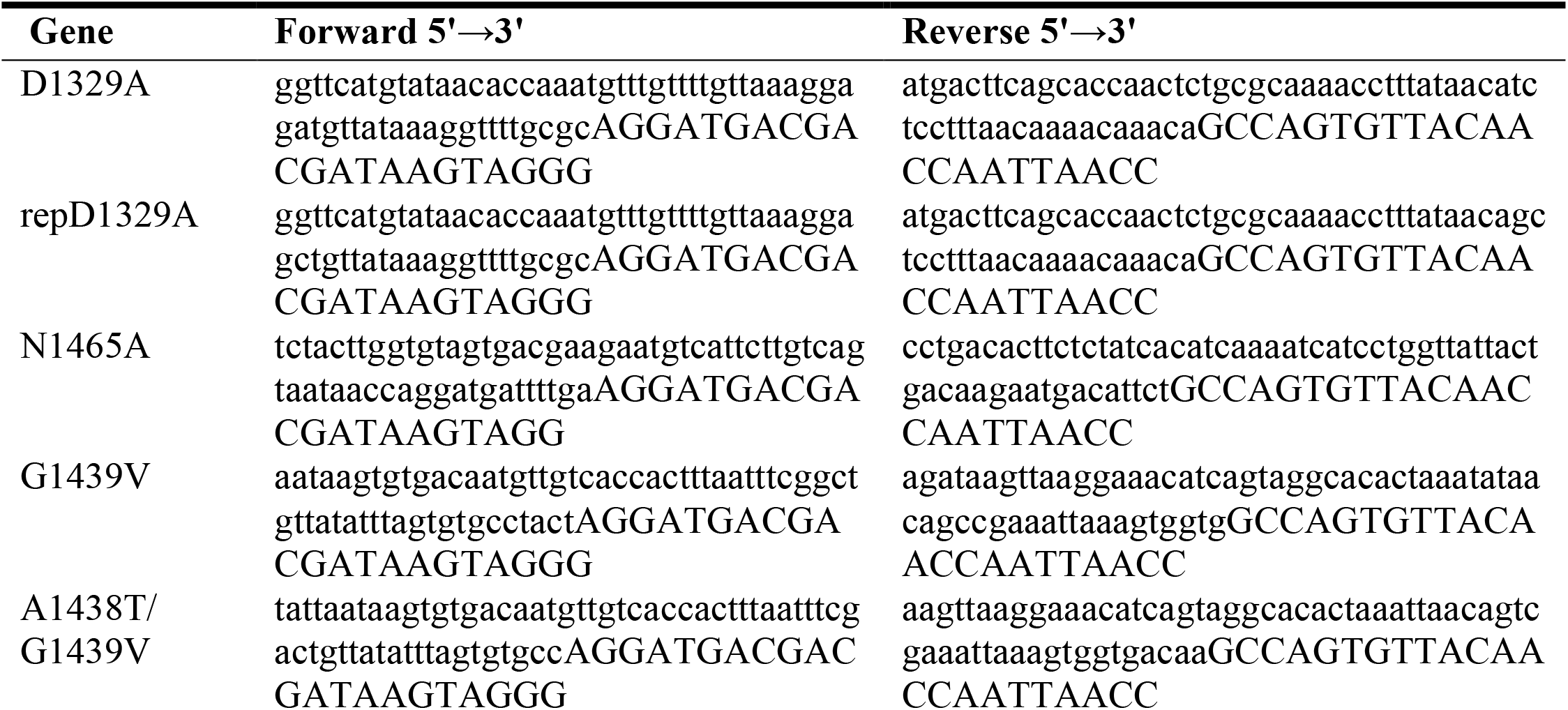
Primers used to create recombinant BACs.

### Reconstitution of recombinant pBAC-JHMV-derived virus

Approximately 1×10^6^ BHK-MVR cells were transfected with approximately 0.5-1 μg of pBAC-JHMV DNA and 1 μg of pcDNA-MHV-N plasmid using Polyjet (SignaGen) as a transfection reagent. New recombinant viruses used in this study were termed D1329A (rJ^IA^-GFP-D1329A), N1465A (rJ^IA^-GFP-N1465), G1439V (rJ^IA^-GFP-G1439V), A1438T/G1439V (rJ^IA^-GFP-A1438T/G1439V), and repD1329 (rJ^IA-^GFP-repD1329). Virus stocks were created by infecting ∼1.5×10^7^ 17Cl-1 cells at an MOI of 0.1 plaque-forming units (PFU)/cell and collecting both the cells and supernatant at 16-20 hpi. The cells were freeze-thawed, and debris was removed prior to collecting virus stocks. Virus stocks were quantified by plaque assay on Hela-MHVR cells and sequenced by collecting infected 17Cl-1 or L929 cells using TRIzol. RNA was isolated and cDNA was prepared using MMLV-reverse transcriptase as per manufacturer’s instructions (Thermo Fisher Scientific). The Mac1 gene sequence was amplified by PCR using the same primers as described above for sequencing BACs, and then resulting PCR products were sequenced by Sanger sequencing using the forward primer. Sequence was analyzed using DNA Star software.

### Virus infection

Cells were infected with recombinant virus at a multiplicity of infection (MOI) of 0.05-0.1 PFU/cell with a 45-60 min adsorption phase, unless otherwise stated. Olaparib (APExBIO – Cat. #A4154) and XAV-939 (APExBIO – Cat. #A1877) were added to cells following the adsorption phase at 10μM. Nicotinomide Riboside Chloride (NR) (Chromadex) was added to cells at 100 μM 4 hr prior to infection and then re-added immediately following the adsorption phase of the infection. Male and female mice, 5 to 8 weeks old (unless otherwise indicated), were anesthetized with ketamine/xylazine and inoculated intranasally with either 3×10^3^ or 3×10^4^ PFU of recombinant virus in a total volume of 12 μl DMEM. To obtain viral RNA from infected animals to sequence the virus following infection, mice were sacrificed, and brain tissue was collected and homogenized in TRIzol (Invitrogen). RNA was isolated and cDNA was prepared using MMLV-reverse transcriptase as per manufacturer’s instructions (Thermo Fisher Scientific). The macrodomain gene sequence was amplified by PCR and sequenced as described above. Sequence was analyzed using DNA Star software.

### Real-time qPCR analysis

RNA was isolated from BMDMs using TRIzol (Invitrogen) and cDNA was prepared as described above. Quantitative real-time PCR (qRT-PCR) was performed on a QuantStudio3 real-time PCR system using PowerUp SYBR Green Master Mix (Thermo Fisher Scientific). Primers used for qPCR are listed in Table 3. Cycle threshold (C_T_) values were normalized to the housekeeping gene hypoxanthine phosphoribosyltransferase (HPRT) by the following equation: C_T_ = C_T(gene of interest)_ - C_T(HPRT)_. Results are shown as a ratio to HPRT calculated as 2^-ΔCT^.

**Table 3.**
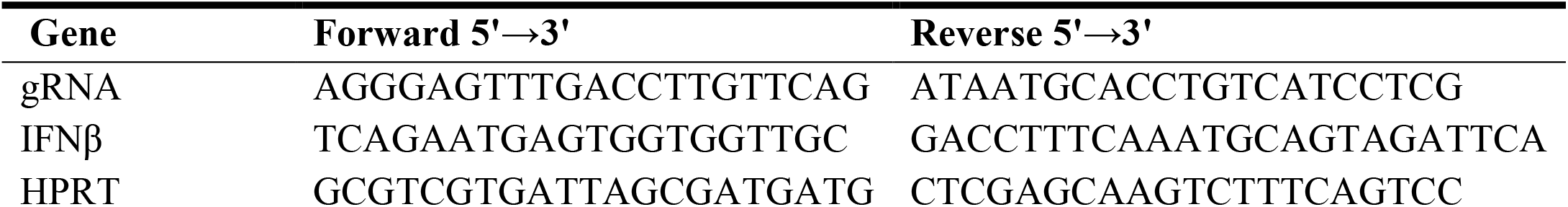
Quantitative real-time PCR primers.

### Immunoblotting

Total cell extracts were lysed in sample buffer containing SDS, protease and phosphatase inhibitors (Roche), β-mercaptoethanol, and a universal nuclease (Fisher Scientific). Proteins were resolved on an SDS polyacrylamide gel, transferred to a polyvinylidene difluoride (PVDF) membrane, hybridized with a primary antibody, reacted with an infrared (IR) dye-conjugated secondary anti-body, visualized using a Li-COR Odyssey Imager (Li-COR), and analyzed using Image Studio software. Primary antibodies used for immunoblotting included anti-MHV N monoclonal antibody (52) and anti-actin monoclonal antibody (clone AC15; Abcam, Inc.). Secondary IR antibodies were purchased from Li-COR.

### Statistical Analysis

All statistical analyses were done using an unpaired two-tailed student’s t-test to assess differences in mean values between groups, and graphs are expressed as geometric mean ± geometric SD (virus titers) or ±SEM (qPCR). The n value represents the number of biologic replicates for each figure. All data was analyzed using GraphPad Prism software.Significant p values are denoted with *p≤0.05, **p≤0.01, or ***p≤0.001.

## ACKNOWLEDGEMENTS

This research was funded by the National Institutes of Health (NIH), grant numbers P20 GM113117, K22AI134993, and R35GM138029, and start-up funds from the University of Kansas to A.R.F. C.M.K. was supported by the NIH Graduate Training at the Biology-Chemistry Interface Grant T32GM132061.

We thank the Davido (University of Kansas), Brenner (City of Hope National Medical Center), and Cohen (Oregon Health Sciences University) labs for insightful discussions on this project; Stanley Perlman, Yousef Alhammad, and Srivatsan Parthisarathy for critical reading of this manuscript; the Perlman lab for reagents; and Chromadex for supplying nicotinamide riboside chloride (NR).

## AUTHOR CONTRIBUTIONS

Conceptualization: ARF, LSV

Data curation: LSV, JJOC, CMK, ARF, DKJ

Formal analysis: LSV, JJOC, ARF, DKJ

Funding acquisition: ARF

Investigation: LSV, JJOC, CMK, ED, NS, PS

Methodology: ARF

Project administration: ARF

Resources: ARF

Supervision: ARF

Validation: LSV, JJOC, CMK, ARF, DKJ

Visualization: LSV, JJOC, CMK, ARF, DKJ

Writing – original draft: LSV, ARF

Writing – review & editing: all authors

